# All-atom MD simulations of the HPV16 capsid

**DOI:** 10.1101/2025.05.22.655575

**Authors:** Santiago Antolínez, Jodi A. Hadden-Perilla

## Abstract

Human papillomavirus (HPV) type 16, one of over 200 known genotypes, is a major cause of anogenital and oropharyngeal cancers worldwide. The L1 protein capsid – which represents the exterior shell of the virus – plays essential roles in host cell adhesion and entry, and is the major antigen used in vaccines that protect against HPV. Here we report an all-atom molecular dynamics (MD) simulation of the intact capsid in explicit solvent, a calculation comprising 16.8 million atoms. The detailed motions and emergent biophysical properties revealed by this simulation provide critical insights into capsid function and immunogenicity, establishing a new platform for rational drug design and antibody engineering.

## Introduction

Human papillomavirus (HPV) is a small (55-60 nm in diameter), non-enveloped DNA virus that infects keratinocytes – flat, plate-like cells that stack to form a barrier tissue called the stratified squamous epithelium. In particular, HPV targets the basal layer of keratinocytes, which represents the deepest tier of actively dividing epithelial cells that can only be accessed through abrasions or microwounds at the surface. More than 200 genotypes of HPV have been identified, with tissue tropisms divided broadly into cutaneous and mucosal categories.

Among mucosal strains, a subset is oncogenic: HPV types 16 and 18 account for the majority of cervical cancers, as well as a rising number of anogenital and oropharyngeal carcinomas in both women and men. Low-risk types, such as HPV 6 and 11, are associated with benign lesions, including genital warts. While most HPV infections are transient and cleared naturally by the immune system, a subset of infections – particularly with high-risk genotypes – persist and can lead to development of cellular abnormalities and malignancies. Although screening tools (such as Pap smears and HPV DNA tests) and clinical interventions enable early detection and treatment of precancerous lesions, there is currently no therapeutic cure for HPV infection itself. The major line of defense against HPV is prophylactic vaccines, which currently protect against up to nine genotypes (6, 11, 16, 18, 31, 33, 45, 52, 58).

Host cell entry begins with virion attachment to heparan sulfate proteoglycans (HSPGs), followed by engagement of secondary receptors, triggering internalization and subsequent endosomal trafficking. The HPV capsid – the exterior shell of the virus – is both the molecular vehicle of infection and the primary antigen used in vaccines. It is composed of 360 copies of the L1 major capsid protein, arranged into 72 pentameric capsomers that are organized with T=7d icosahedral symmetry (**Figure 1**). Each L1 monomer adopts a classical *β*-strand jellyroll fold and contributes a flexible C-terminal arm that interlocks with adjacent capsomers, forming a network of linkers essential for capsid stability (**Figure 1a**). Twelve of these capsomers occupy the icosahedral vertices and are classified as pentavalent, while the remaining sixty occupy hexavalent (pseudo-sixfold) positions, despite their fivefold symmetry (**Figure 1b**). Each pentavalent capsomer and the five hexavalent capsomers surrounding it form a structural subunit called a “starfish” (a pentamer of pentamers), and the architecture of the complete capsid can be conceptualized as twelve tethered “starfish” (**Figure 1c**). The L1 protein alone is sufficient to drive self-assembly into virus-like particles (VLPs, *e*.*g*. the empty L1 capsid), which form the basis of the three FDA-approved vaccines in the United States: Cervarix (bivalent), Gardasil (quadrivalent), and Gardasil 9 (nonavalent).

**Figure 1:**
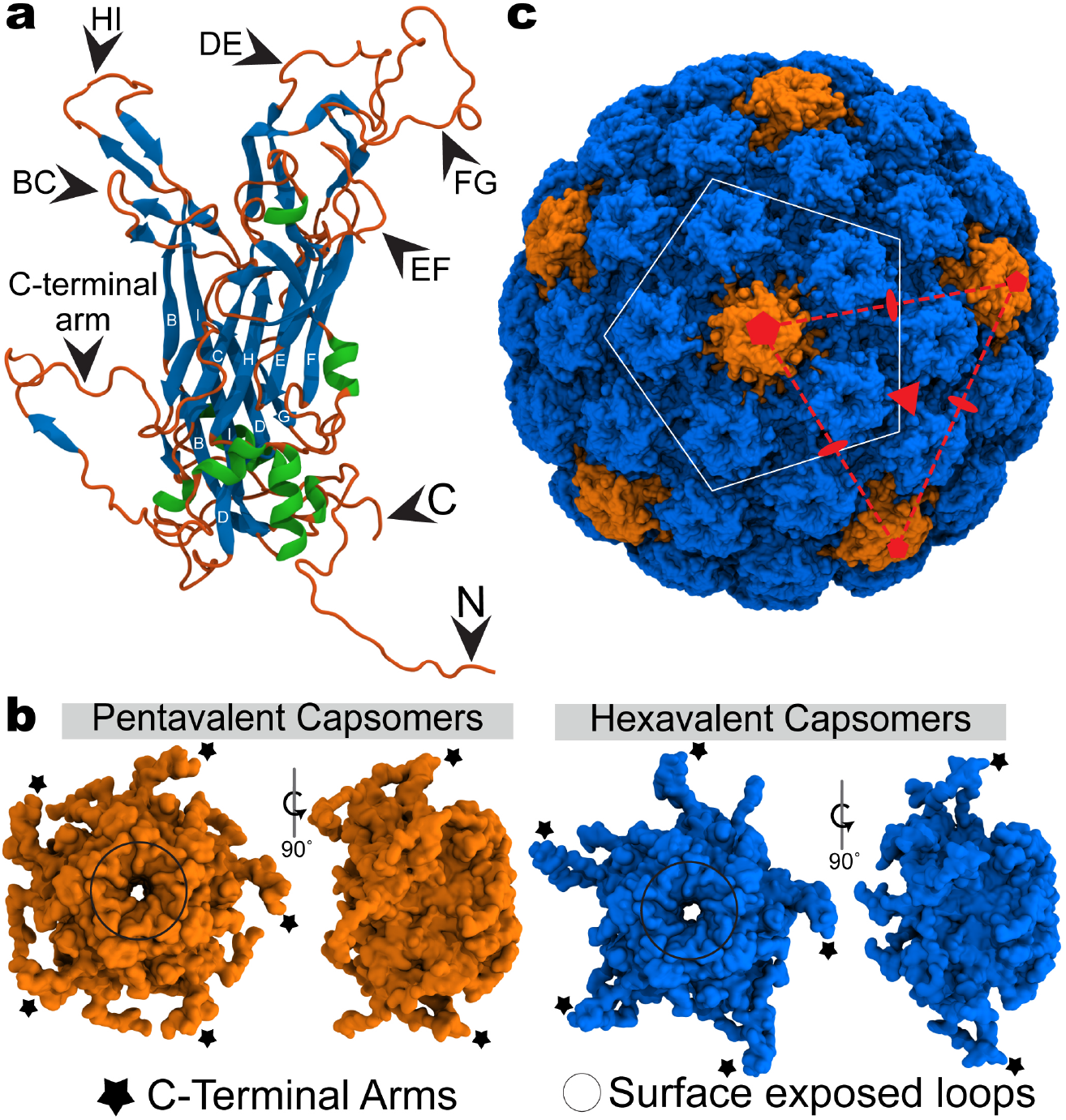
**(a)** L1 major capsid protein folds into a classical *β*-jelly roll, composed of two antiparallel sheets, BIDG and CHEF, connected by five surface exposed loops (BC, DE, EF, FG, and HI). **(b)** Five L1 subunits assemble into pentameric capsomers. The twelve capsomers on the icosahedral vertices are pentavalent (five neighbors, orange), while the remaining 60 are hexavalent (six neighbors, blue). Capsomers are interlocked via the L1 C-terminal arm. **(c)** HPV virus-like particle, an empty capsid composed of 360 copies of L1 arranged into 72 capsomers organized with T=7d icosahedral symmetry. The “starfish” subunit – a pentamer of pentamers, of which there are 12 per capsid – is denoted in white.

High-resolution cryo-electron microscopy (cryo-EM) has provided atomic models of the HPV capsid, most notably for type 16, revealing the global organization of capsomers and localized features of the L1 *β*-strand jellyroll. However, the capsid exhibits a high degree of conformational plasticity, particularly in the flexible loops and termini that are often unresolved in experimental reconstructions. Global motions of the capsid further contribute to resolution averaging, making certain functional regions structurally elusive. Recent subparticle refinement methods have partially overcome these limitations, but many questions remain about the dynamic behavior of the capsid and how its motion contributes to infection. Further, the atomic details of L2 minor capsid protein incorporation into the VLP represents a major open question in HPV biology. Though typically present at lower copy number (*∼*12–72 per virion), L2 binds viral DNA and participates in genome encapsidation, nuclear import, endosomal escape. Experimental evidence supports a model in which L2 inserts through the center of L1 capsomers, with its C-terminus extending into the lumen to contact genome and its N-terminus exposed on the exterior to mediate intracellular trafficking. However, structural characterization of L2 within the native capsid remains wholly incomplete.

Molecular dynamics (MD) simulations provide a complementary approach for interrogating viral capsids at atomic resolution and on biologically relevant timescales. Here, we present an all-atom MD simulation of the intact HPV type 16 capsid in explicit solvent, comprising 16.8 million atoms. This simulation – the largest yet reported for an icosahedral capsid from a human-infective virus – enables unprecedented characterization of HPV’s conformational dynamics, providing critical insights into the capsid’s function during infection. Beyond guiding the interpretation of experimental results to bridge long-standing knowledge gaps, the detailed conformational ensemble afforded by this dataset establishes a new platform for rational drug and immunogen design that accounts for the capsid’s innate plasticity.

## Results

### Dynamics of capsid-incorporated L1

Root-mean-squared fluctuation (RMSF) of L1 indicates that the *β*-strands of the jellyroll – essentially the core of the protein – are relatively stationary (**Figure 2a-b, top**). By comparison, the N-terminal tail, surface exposed loops, and C-terminal arm are quite mobile, with their C*α* coordinates varying up to 3 Å per residue locally. This uncertainty in position accounts for the absence of these L1 regions from all but the most recent and highest-resolution reconstructions. The C-terminal tail is by far the most fluid portion of L1, consistent with this peptide – which is thought to be disordered – being unresolved in all experimentally derived models obtained to date. Regardless of spatial fluctuation, the secondary structure captured by cryo-EM (PDB 7KZF^1^) remained highly conserved during MD, with 80% of residues exhibiting the expected local folding pattern in over 90% of sampled conformers (**Figure 2b, bottom**).

**Figure 2:**
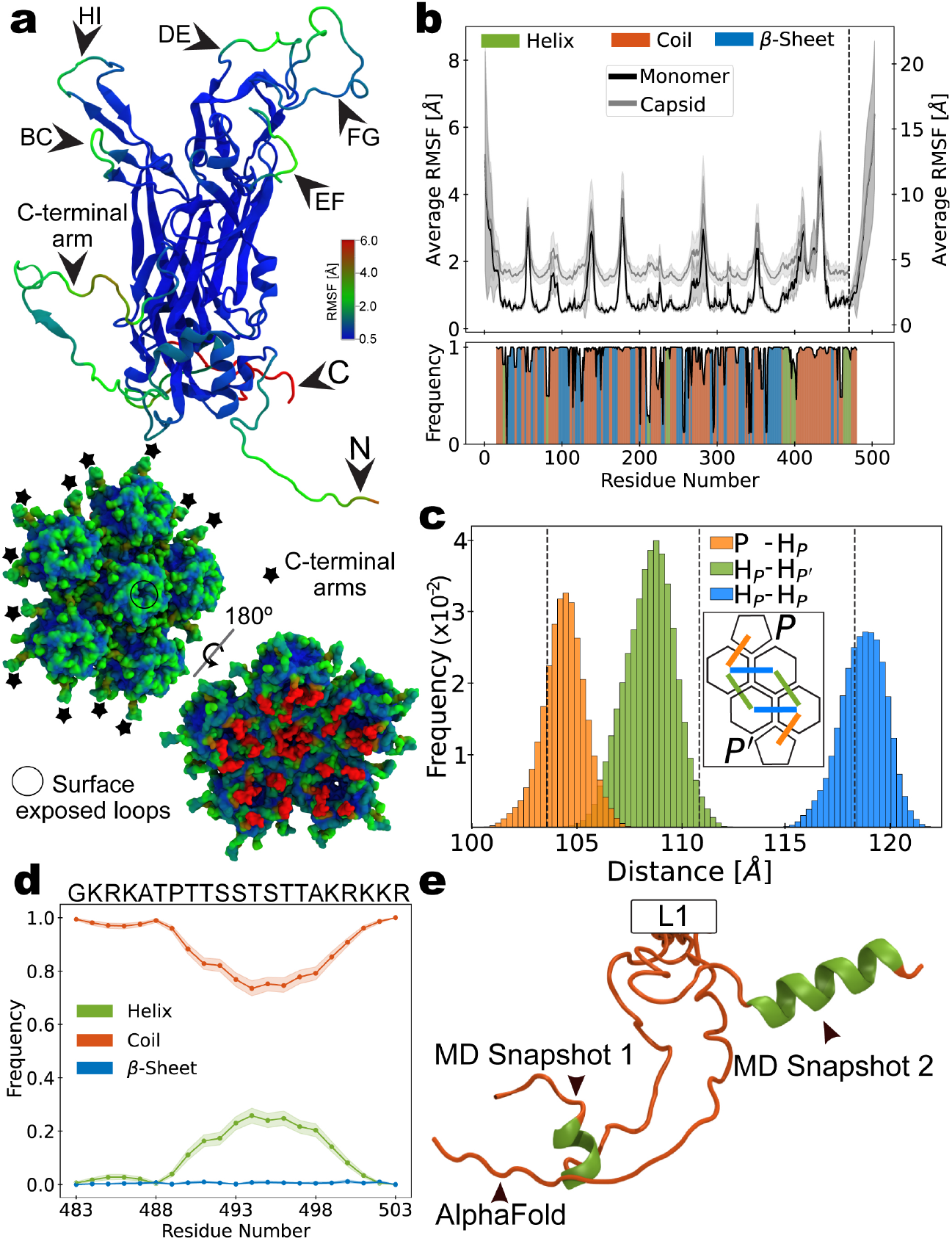
**(a)** Average C_*α*_ RMSF projected onto L1 monomer (top) and “starfish” capsid fragment (bottom). **(b)** Average C_*α*_ RMSF based on internal alignment of L1 extracted from the capsid (local dynamics *[e*.*g*., *panel a]*, black) and alignment of the intact capsid (global dynamics, gray). Dashed line separates sections of plot using left and right x-axis scales (top). Conservation of secondary structure reported by cryo-EM (bottom, PDB 7KZF^1^). **(c)** Distributions of inter-capsomeric distances, measured between geometric centers of adjacent capsomers. Inset diagram illustrates quasi-equivalent relationships between pentavalent (P) and hexavalent (H) capsomers in the T=7d icosahedral lattice. Dashed lines indicate analogous distances from cryo-EM (PDB 7KZF^1^). **(d)** Frequency of secondary structure observed for C-terminal tails, aggregated over 9 million conformational samples. **(e)** Comparison of AlphaFold^2^ model with two conformers extracted from MD: minimal helix (snaphot 1, one turn, 9.5% of conformational ensemble, observed in 184 separate copies of L1) and a maximal helix (three turns, 0.5% of conformational ensemble, observed in 26 separate copies of L1).

### Spontaneous ordering of C-terminal tail

Given what has been previously inferred about their nature, the C-terminal tails were modeled into the MD initial coordinates as disordered peptides. AlphaFold^2^ predicts a disordered configuration for this portion of L1, and MD simulations of the isolated peptide in implicit solvent suggested a preference to remain in a disordered state. However, during MD simulations of the intact capsid in explicit solvent, select C-terminal tails were observed to adopt a helical secondary structure (**Figure 2d**). The helix spanned up to three turns over residues 489-501, with the central turn being most common (**Figure 2e**). Across all MD-derived conformers collected for all 360 copies of L1 (over 240 µs of cumulative sampling)), the peptide exhibited a 0.25 probability to contain a helical element. Upstream residues 473-488 may serve as a linker connecting the helix to the *β*-strand jellyroll core of the protein, allowing it to alter its orientation as needed for function. Possibly, the C-terminal helix is stabilized by the presence of incorporated L2 or encapsidated DNA. In that case, it could be the flexibility of the purported linker – rather than disorder of the entire C-terminal tail – that accounts for the lack of resolution for this portion of L1 in experimentally-derived models.

### Global dynamics of intact L1 capsid

Beyond the internal motions of L1, the capsid assembly exhibits remarkable global dynamics. When L1 is considered in the context of the complete assembly, the spatial translations indicative of global dynamics manifest as an up-shift in the C*α* RMSF profile (**Figure 2b, top**). Because the HPV capsid is composed entirely of pentameric capsomers, 60 of these serve as obligate hexamers in the T=7d icosahedral lattice. As a consequence, these groups of five subunits each occupy a space theoretically large enough to accommodate six subunits, leading to capsomer mobility along the surface. MD simulations reveal the variation in relative capsomer positions, indicating that the distance between capsomers can fluctuate by up to 6 Å. Consistent with cryo-EM studies of the apo-form capsid,^1^ hexavalent capsomers were found, on average, to sit closer to pentavalent capsomers than to other hexavalent capsomers (**Figure 2c**). Given that HSPG host receptors and their analogs (*e*.*g*., heparan sulfate and heparin), as well as several HPV-targeting antibodies (*e*.*g*. U4), are known to specifically recognize the canyon between pentavalent and hexavalent capsomers,^3,4^ the observation that this inter-capsomeric distance is smallest, with the most narrow canyon, is likely relevant to the capsid’s function in cell adhesion, as well as immunogenicity.

### Morphology and interior compartment

Changes in inter-capsomer positioning suggest shifts in size and morphology. Indeed, distributions of capsid diameters for MD-derived conformers indicate subtle expansion of the shell relative to the cryo-EM structure (**Figure 3a**). Diameter quantification depends on symmetry axis, reflecting the icosahedral rather than spherical shape of the assembly. Increases in capsid dimensions during equilibration have been reported for MD simulations of other capsids,^6,7^ and have been attributed to relaxation from low-energy crystal-packed or vitrified states to native conditions at physiological temperature. With respect to the capsid lumen, the volume of interior space enlarged by 1.5% relative to the value reported by cryo-EM (**Figure 3b**), consistent with the scale of expansion observed for diameter. However, owing to small adjustments in capsid morphology, lumen enlargement is not uniformly distributed. MD simulations show that, as the capsid relaxes to native conditions, its sphericity slightly decreases (**Figure 3c**). Reduced sphericity arises from minute protrusion of the icosahedral vertices, concomitant with flattening of the icosahedral faces (**Figure 3d-e**). These changes align with the expansion of spacing between pentavalent and hexavalent capsomers within “starfish” centered at the fivefold axes, and the compaction of spacing between hexavalent capsomers where neighboring “starfish” connect at the threefold axes (**Figure 2c**). Relative to the cryo-EM structure, the average faceting angle (*φ*_5_ *− φ*_3_) increased by 1.9 *±* 0.1^*°*^ during MD (**Figure 3e**). Given that a perfect sphere and truncated icosahedron have sphericity of and 0.92, respectively, the HPV capsid remains highly sphere-like in shape within a physiological environment. Although these underlying structural adjustments appear negligible on the scale of the intact capsid, distance shifts of only a few Ångströms remain significant at the atomistic level, and can mean the difference in a key chemical contact being within range for interaction or not, with potential consequences for biological function.

**Figure 3:**
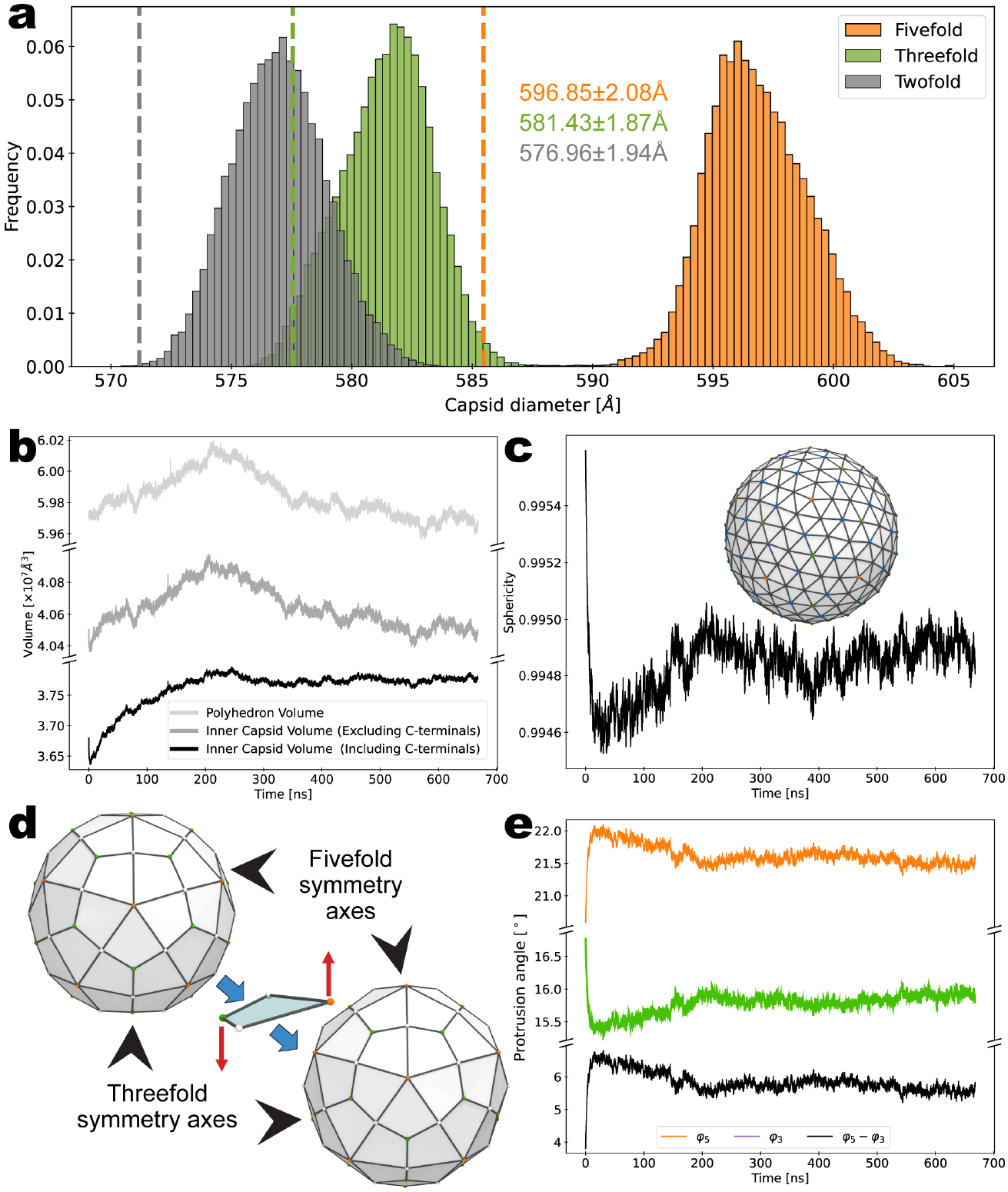
(**a**) Distributions of capsid diameters measured as the difference between maximum and minimum atomic coordinates along icosahedral symmetry axes. **(b)** Capsid shell volume represents the contained space of a geodesic polyhedron of 420 triangular faces fitted to the protein assembly based on its symmetry vertices (panel b inset). Inner volume approximates the dimensions of the capsid lumen, as detected by VMD’s *measure volInterior*^5^ functionality. **(c)** Capsid sphericity, a relationship between shell volume and surface area [Φ = *π*^1*/*3^(6*V*)^2*/*3^*/SA*], estimated based on the same geodesic polyhedron (inset). **(d)** Capsid faceting occurs when the fivefold vertices (orange) protrude and the threefold vertices (green) flatten, causing the global morphology to approach that of a regular icosahedron with distinct triangular faces. **(e)** Faceting angles indicate the orientation of the capsid asymmetric unit relative to the icosahedral symmetry axes. The extent to which the orientation causes fivefold protrusion and threefold flattening, captured by *φ*_5_ − *φ*_3_ (black), quantifies the degree of faceting. More pronounced faceting is correlated with reduced sphericity.

### Interactions with solvent environment

Previous MD simulation studies on intact virus capsids have demonstrated that these biological containers exhibit complex interactions with their solvent environment, and the HPV capsid is no exception. The capsid exchanges water molecules across its surface at a rate of 6,400 ns^*−*1^ (**Figure 4a**). Equal rates for inward versus outward transport of water indicate equilibrium between the lumen and exterior. Given this rate of exchange, the empty L1 capsid – equivalent to a monovalent vaccine particle – completely recycles its internal solvent every *∼*200 ns. With respect to salt, the capsid behaves as a selectively permeable container, exchanging Cl^*−*^ ions at a rate five times faster than Na^+^ ions (approximately 5 versus 1 ions per nanosecond, respectively). The relative number of ions included in the MD simulation does not account for this difference, with the system containing a 1.3/1.0 ratio of Cl^*−*^/Na^+^ ions. Again, equal rates for inward versus outward transport of ions indicates equilibrium with the surroundings, and highlights the capsid’s ability to regulate its internal salinity to maintain homeostasis. While the bulk concentration of NaCl was set to 150 mM, the bulk concentration of internal solvent was found to be 177*±*3 mM NaCl, revealing higher ionic strength within the capsid lumen. Inside the empty L1 capsid, this likely provides some electrostatic compensation for the absence of DNA and the L2 C-terminus. The capsid’s preference to transport negatively, rather than positively charged species may relate to key biological functions, such the import of electronegative cofactors or genome uncoating.

**Figure 4:**
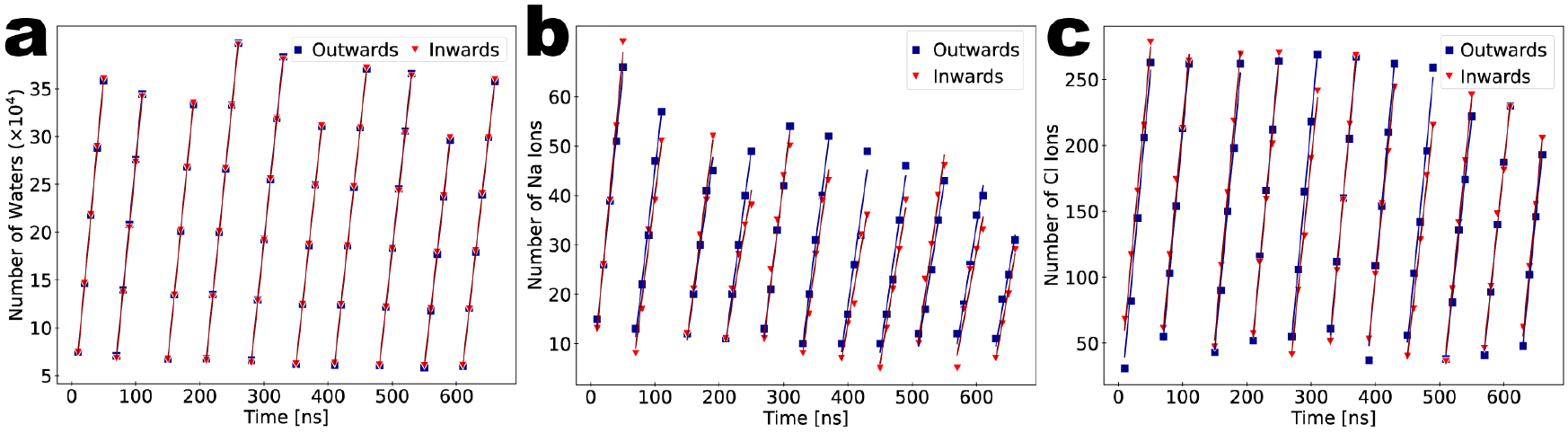
Rates of solvent exchange across the capsid surface, moving inward from exterior to lumen (red) and outward from lumen to exterior (blue). **(a)** Water molecules: inward (6,400 *±* 410 ns^*−*1^) and outward (6,410 *±* 410 ns^*−*1^). **(b)** Sodium ions: inward (0.91 *±* 0.21 ns^*−*1^) and outward (0.96 *±* 0.16 ns^*−*1^). **(c)** Chloride ions: inward (5.01 *±* 0.35 ns^*−*1^) and outward (5.11 *±* 0.31 ns^*−*1^). Equal inward/outward rates indicate equilibrium.

### Topology and dynamics of capsid pores

Equilibrium with the surrounding solvent environment is mediated by the capsid’s pores, located at the centers of capsomers. Solvent exchange was found to occur through both pentavalent and hexavalent capsomer pores, whose channels exhibit a characteristic hourglass profile (**Figure 5a**). While the dimensions of the channels are, on average, similar for the two types of capsomers, the pentavalent pores tend to have a smaller bottleneck (**Figure 5b**). MD simulations indicate, consistent with cryo-EM, that their minimum radius is 3.71 *±* 0.56 Å (**Figure 5c**). However, the hexavalent pores were observed to be statistically less constricted, with a minimum radius of 4.47 *±* 0.43 Å (**Figure 5c**). Fluctuations of hydrophobic residues Phe256 and Met299 control the size of the bottleneck, allowing passage of solvent and small molecules up to 11 Å across, assuming pores are not occluded by the incorporation of L2. The combined cross-section of the most narrow pore apertures represents the cumulative area connecting the capsid lumen with the exterior. The total cross-sectional area is dominated by the nature of hexavalent capsomers, which are five times more numerous than their pentavalent counterparts in the complete T=7d structure (**Figure 5d**).

**Figure 5:**
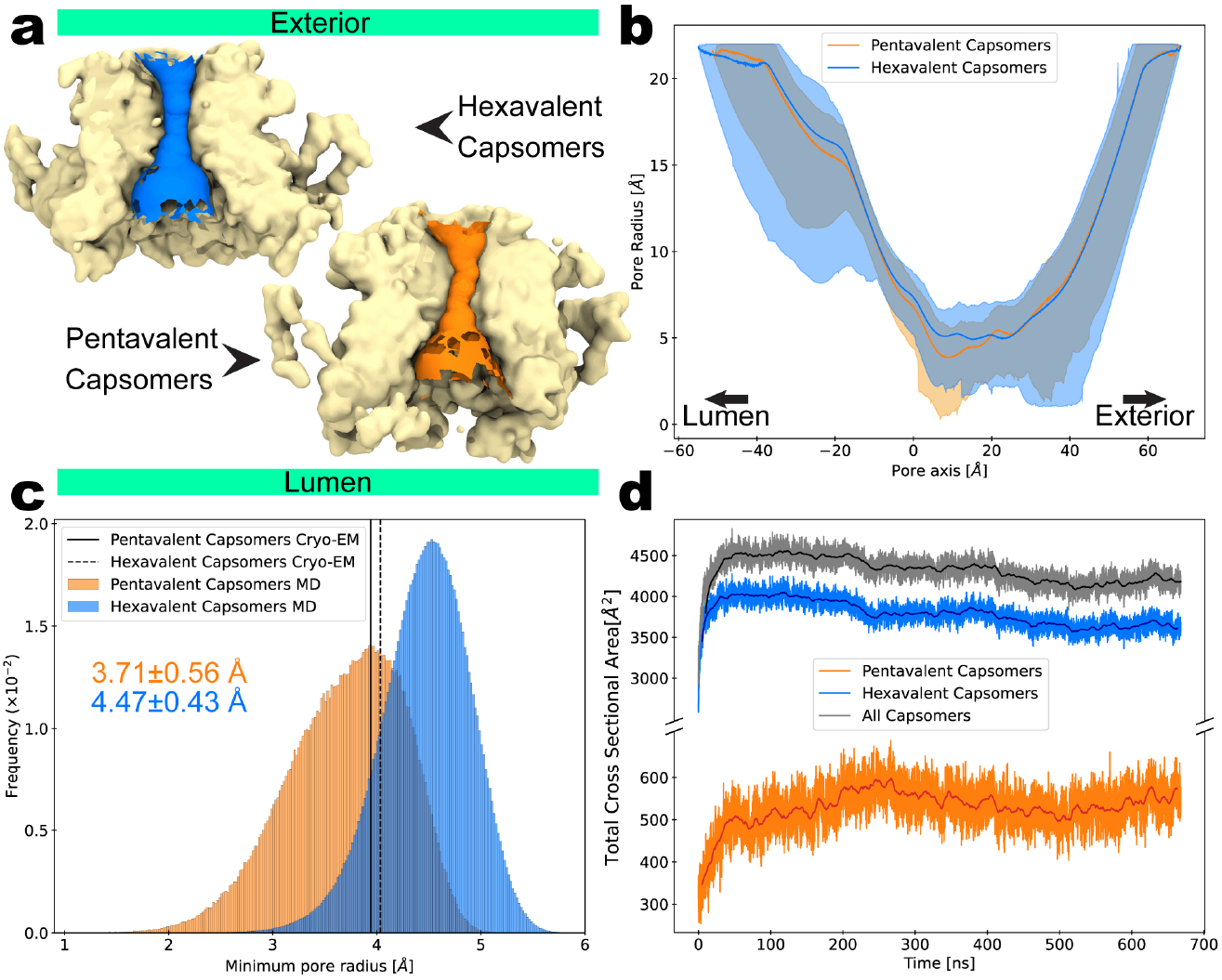
**(a)** Capsomer cross-sections showing the hourglass shape of the pores that connect the capsid lumen with the exterior, as calculated by HOLE.^8^ **(b)** Average radial profiles of the pores along their channel axes, where shading indicates the minimum/maximum values observed. **(c)** Distributions of minimum pore radii. Solid and dashed lines indicate analogous values from cryo-EM (PDB 7KZF^1^): 3.94 Å for pentavalent and 4.03 Å for hexavalent capsomers, respectively. **(d)** Total cross-sectional area associated with minimum pore radii.

## Discussion

Cryo-EM studies of the HPV capsid suggest that it is highly dynamic, with mobile capsomers connected by flexible linkers – namely, the C-terminal arms of the L1 protein. MD simulations confirm this plasticity, revealing internal flexibility within L1 subunits, variability in inter-capsomer positioning, and global fluctuations in capsid shape and diameter. These motions clearly contribute to resolution blurring historically observed in cryo-EM maps, particularly in the flexible loop and terminal regions. The success of more recent sub-particle refinements stems from the omission of global motions and suppression of asymmetries that convolute datasets. MD simulations show that, in the case of HPV, focus on local conformational space reduces uncertainty in atomic coordinates by *∼*1 Å (**Figure 2b, top**).

Prior work has reported mature HPV capsid diameters typically in the range of 55–60 nm. Capsid diameter was found to vary depending on the axis of measurement for MD-derived conformers, however, inconsistent particle orientations do not account for size variation observed in microscopy studies.^1,4,9^ Consistent with MD data, cryo-EM classes with discrete diameters exhibit Gaussian distributions,^1^ and this heterogeneity is recognized as a limiting factor for resolution in HPV structures. Initial coordinates for the simulation were taken from PDB 7KZF,^1^ which has a diameter of 58.5 nm when measured as the difference between maximum and minimum atomic positions along the fivefold symmetry axis. The reconstruction was based on a population of particles whose diameters spanned *±*2*σ* ≈ 0.92 nm about the value of the icosahedrally averaged map, suggesting a spread of 57.6–59.4 nm. The corresponding diameters predicted for the model by MD – with an average of 59.7 nm and spread of 59.3–60.1 nm (**Figure 3a**) – represent a similar size distribution. This result demonstrates that vitrification during cryo-EM sample preparation does not meaningfully underestimate the capsid dimensions expected at native physiological conditions.

Mature HPV capsids are stabilized by inter-capsomeric disulfide linkages formed between neighboring L1 subunits (*e*.*g*., Cys428 and Cys175), which arise during transit through the oxidizing environment of the upper epithelium. Following particle assembly, immature virions present as genome-containing, yet loosely-connected procapsids with diameters *>*60 nm.^10^ Accumulation of disulfide bonds is associated with viral maturation, in which the cross-links tighten the capsid lattice, leading to anisotropic structural contraction. Particle diameters are reduced by 12%, 5%, and 3% along the fivefold, threefold, and twofold symmetry axes, respectively,^10^ leading to more spherical morphologies. Because the MD simulation model incorporated all 360 possible disulfides, the predicted diameters may reflect the lower end of the size range for capsids at native physiological conditions. Particles that contain inconsistent or asymmetric disulfide bonding patterns are likely to exhibit expanded diameters associated with incomplete maturation. Although *in vitro* assays show that maturation is not strictly a prerequisite for host infectivity, the compaction and cross-linking process – which proceeds over several hours – markedly increases virion stability.^10^

Measurements on MD-derived conformers indicate that the L1 capsid lumen encompasses a volume of *∼*37,700 nm^3^ at physiological conditions (**Figure 3b**), representing the space available for the 8,000 base pair genome and other packaged cargo. The double-stranded HPV DNA is spooled onto host-acquired histone complexes, producing 30 viral nucleosomes per particle.^11^ Various schemes have been proposed to describe the organization of nucleosomes into chromatin, including the zigzag fiber model, which is supported primarily by *in vitro* evidence.^12–14^ Based on this two-start helical arrangement of 6.5 nucleosomes per 11 nm turn,^15^ the HPV genome would require a cylindrical volume of *∼*35,800 nm^3^ – highly complementary to the interior compartment characterized by MD simulations. Given that monovalent salt has been shown to induce chromatin compaction,^16^ the L1 capsid’s ability to maintain elevated internal salinity reletave to its external environment appears to promote efficient packaging. It may be that the process of particle compaction during virion maturation^10^ is also related to late-stage genome compaction, with disulfide cross-links gradually locking DNA into an optimally sized homeostatic compartment.

Nevertheless, accumulating evidence implicates liquid-liquid phase-separation (LLPS) as an important driver of chromatin organization *in vivo*, supporting a framework in which nucleosome association is fluid and lacks the long-range order imposed by fiber models.^17^ HPV capsid assembly proceeds in the complete absence of DNA, such that the genome does not function as a space-filling structural scaffold for particle formation. Beyond being entropically favorable, a dynamic arrangement of DNA could more readily adapt to the constraints of an auto-assembling container during packaging. Moreover, an irregular and globally disordered genome structure, perhaps unique across particles, is consistent with the lack of ordered density corresponding to packaged DNA in symmerized cryo-EM reconstructions of pseudovirions. A static model of the L1 capsid containing a fluid rather than fiber like association of nucleosomes was recently used to explain the effects of histone modifications.^18^

MD simulations reveal a range of conformations adopted by the L1 C-terminal tail within the context of the intact capsid, providing structural insight into a region that has consistently eluded experimental resolution. In particular, residues 489–500 exhibited transient *α*-helical character, with up to 25% probability of forming secondary structure. The flexibility of this domain, along with the loop connecting it to the L1 core, likely accounts for its persistent absence in high-resolution cryo-EM reconstructions. Given that the last nine residues of the L1 C-terminal tail have been implicated in genome packaging (*e*.*g*. 495–503 of sequence STTAKRKKR, **Figure 2d-e**),^19^ the predicted helical elements are likely relevant to binding DNA. Biologically, the tail is multifunctional. Beyond a role in genome interactions, it can also drive nuclear localization of L1 during early infection or late-stage viral assembly.^20,21^ Further, the extreme C-terminus may participate in binding to cell surface HSPGs, which are critical for initial host attachment.^22^ These diverse functions suggest that the C-terminal tail alternates between lumen-facing and externally-exposed states, depending on stage of the viral life cycle. This dynamic behavior is consistent with the presence of both disordered and transiently folded conformers and reflects a broader trend observed in other viral capsid disordered domains that mediate both genome interactions and cellular signaling.

The *α*-helix observed in simulations of the L1 C-terminal tail consisted of up the three turns of sequence TTSSTSTTAKRK (**Figure 2d-e**). Although lacking enrichment with arginine and lysine residues typical of canonical DNA-binding helices in all but the final turn, the upstream portion of the helix may still contribute functionally to genome packaging. Structured DNA-binding helices are often stabilized by hydrogen bonds and shape complementarity within the DNA major groove – a feature partially recapitulated by the helical geometry observed here. However, the T/S-rich composition of this L1 helix is perhaps more consistent with a role in structural support, protein–protein interactions, or post-translational control rather than direct DNA contact. The abundance of threonine and serine residues introduces potential sites for phosphorylation, suggesting that this region may also function as a regulatory domain that modulates L1 activity, trafficking, or interactions with viral or host partners. It is conceivable that L1 provides structural positioning for L2, which contains basic DNA-binding motifs and serves as the primary DNA recognition element. Collectively, these data support a model in which the L1 C-terminal tail could contact DNA directly or act as a dynamic scaffold – functionally plastic and responsive to both structural and regulatory cues during the HPV life cycle, in roles both internal and external to the capsid.

## Methods

### Molecular modeling

Initial coordinates for the L1 protein capsid were taken from a previously reported cryo-EM structure (PDB: 7KZF).^1^ Unresolved N-terminal residues (chain A:1-11, chain B:1-15, chain E:1-2) were modeled based on homology to chain C. Unresolved residues upstream of the C-terminal tails (chain B:481-482, chain C/D:481-485, chain E/F:482-485) were modeled based on homology to chain A. C-terminal residues 486-503 for all six chains were modeled based on conformers generated by *ab inio* MD of the isolated peptide in Generalized Born implicit solvent:^23^ Starting from an extended structure, simulations were run for 7 µs at 310 K. From the resulting ensemble, disordered conformers that produced the least number of clashes with the experimentally-derived portion of the protein were selected to complete each chain. Atomic overlap of residues in the cryo-EM structure (chain A/F:87-90), as well as any close contacts introduced by the modeled C-terminal tails, were addressed by energy minimization in Generalized Born implicit solvent.^23^ A model of the complete, symmeterized capsid was constructed by applying the provided^1^ translations and rotations to the asymmetric unit. PDB2PQR^24^ was used to add hydrogen atoms appropriate for pH 7.0 with the propKa algorithm,^25^ considering the interfaces of each capsid-incorporated chain.

### System construction

Local ions were placed around the capsid using the Cionize plugin in VMD 1.9.4,^26^ and the model was immersed in a truncated octahedron of TIP3P water^27^ containing 150 mM NaCl. The simulation box was defined such that the distance between hexagonal faces of the truncated octahedron was 30 Å greater than the capsid diameter. Following 136 ns of unrestrained dynamics, to accommodate capsid expansion and box condensation that occurred during relaxation, the simulation box size was increased by placing an additional layer of solvent around the system: A larger truncated octahedron of TIP3P water^27^ containing 150 mM NaCl was generated, such that the distance between hexagonal faces was 30 Å greater than the solvated capsid system. This pure solvent box was equilibrated for 7 ns at 310 K to converge its density, then the outer solvent layer – in excess of the solvated capsid system dimensions – was extracted and merged with the solvated capsid system. Any atomic overlaps at the union boundary were addressed by removing solvent molecules from the added layer, so as to maintain all constituents of the original system. Instantaneous velocities for all atoms were retained in the merged system upon restarting dynamics. The final simulation system for the HPV type 16 L1 protein capsid comprised 16.8 million atoms.

### Molecular dynamics simulations

MD simulations were performed with NAMD3^28^ on the TACC Frontera supercomputer. Protein parameters were taken from the AMBERff-in-NAMD^29^ implementation of the ff14SB force field.^30^ Energy minimization with the conjugate gradient algorithm was carried out in two stages: 10,000 cycles to relax the solvent around the protein, followed by 10,000 cycles to relax the solvated protein sidechains, applying harmonic positional restraints (5 kcalmol ^*−*1^Å ^*−*2^) to the backbone. The temperature of the system was gradually raised from 60 K to 310 K over a period of 5 ns, then harmonic restraints were released over a subsequent 5 ns. Dynamics were explored in the isothermic-isobaric (NPT) ensemble, using a time step of 2 fs. Covalent bonds between hydrogens and heavy atoms were constrained using the SHAKE and SETTLE algorithms for solute and solvent, respectively. Temperature was maintained at 310 K with the Langevin thermostat, using a damping coefficient of 1.0 ps^*−*1^. Pressure was maintained at 1 bar with the Nose-Hoover Langevin piston barostat, using an oscillation time scale of 2000 fs and damping time scale of 1000 fs. A cutoff of 8 Å was used to distinguish between long and short range non-bonded interactions. Long range electrostatics were calculated every other time step with the particle mesh Ewald (PME) method.

### Trajectory analysis

MD trajectory analysis and visualization was carried out with VMD 1.9.4.^26^ Determination of capsid inner volume with VMD’s *measure volInterior*^5^ functionality used the fixed-boundary implementation with molecular surface parameters: Radius Scale = 2.5 Å, Isovalue = 0.5, Grid Spacing = 1.0 Å, and N_*rays*_ = 32. Characterization of capsid shell volume and related morphological properties used polyhedra fitted to the icosahedral structure as described in Refs.^7,31^ Solvent exchange rates we calculated as described in Ref.^7^ Capsid pores were mapped using HOLE.^8^

## Generative AI declaration

During the preparation of this work, the authors used ChatGPT version 4o to improve clarity of language and readability. After using this tool, the authors reviewed and edited the material as needed and take full responsibility for the content of this manuscript.

## Acknowledgments

This work was funded by the NIH through the Center of Biomedical Research Excellence (Discovery of Chemical Probes and Therapeutic Leads) at the University of Delaware, via an Administrative Supplement for Research on Women’s Health in the IDeA States, award 3P20-GM-104316-09S1 to J.A.H.-P. This work was also supported by the Delaware Advanced Research Workforce and Innovation Network (DARWIN), funded by NSF award OAC-1919839, and the BioStore resource, made possible by NIH through the Delaware IDeA Network of Biomedical Research Excellence, awards P20-GM-103446 and S10-OD-028725. Computer time on Frontera at the Texas Advanced Computing Center at the University of Texas at Austin was provided by allocation MCB-23072. The authors are grateful to Dr. David J. Hardy of the Theoretical and Computational Biophysics Group at the University of Illinois at Urbana-Champaign for deployment of NAMD 3.0beta5 on Frontera.

